# Prediction of Golgi Polarity in Collectively Migrating Epithelial Cells Using Graph Neural Network

**DOI:** 10.1101/2022.08.04.502886

**Authors:** Purnati Khuntia, Tamal Das

**Affiliations:** Tata Institute of Fundamental Research Hyderabad (TIFR-H), Hyderabad – 500 046, India

**Keywords:** Collective cell migration, Golgi apparatus, Cell polarity, Graph neural network, Deep learning

## Abstract

In the stationary epithelium, the Golgi apparatus assumes an apical position, above the cell nucleus. However, during wound healing and morphogenesis, as the epithelial cells starts migrating, it relocalizes closer to the basal plane. On this plane, the position of Golgi with respect to the cell nucleus defines the organizational polarity of a migrating epithelial cell, which is crucial for an efficient collective migration. Yet, factors influencing the Golgi polarity remain elusive. Here we constructed a graph neural network-based deep learning model to systematically analyze the dependency of Golgi polarity on multiple geometric and physical factors. In spite of the complexity of a migrating epithelial monolayer, our simple model was able to predict the Golgi polarity with 75% accuracy. Moreover, the model predicted that Golgi polarity predominantly correlates with the orientation of maximum principal stress. Finally, we found that this correlation operates locally since progressive coarsening of the stress field over multiple cell-lengths reduced the stress polarity-Golgi polarity correlation as well as the predictive accuracy of the neural network model. Taken together, our results demonstrated that graph neural networks could be a powerful tool towards understanding how different physical factors influence collective cell migration. They also highlighted a previously unknown role of physical cues in defining the intracellular organization.

## INTRODUCTION

Collective cell migration describes the coordinated movement of a group of cells, where cellular motions and intracellular structures are correlated over a distance that is much larger than the diameter of an individual cell (1, 2). It can occur in form of epithelial sheets (3), cell clusters, or strands of cells (4, 5). Collective cell migration is critical to tissue remodeling during development (6, 7), metastatic invasion of cancer cells (8), and wound closure (9, 10). In wound closure, the directed collective migration of epithelial cells marks the beginning of the process. Consequently, at the onset of collective migration, stationary epithelial cells undergo a drastic intracellular reorganization to acquire the required migratory phenotype (1, 11). For example, in the stationary epithelium, the cell nucleus resides close to the basal plane, and the Golgi apparatus stays on top of the nucleus - towards the apical plane of the cell - establishing the apico-basal polarity axis (11). However, a change in Golgi position from the top of the nucleus towards the basal plane precedes the cell migration. This brings about the well-known change from apico-basal polarity, which is characteristic of the stationary cells, to front-to-back polarity, which is characteristic of the migrating cells (1). In a recent study, we described the nature of this apical-to-basal Golgi relocalization (11), where we found that Golgi undergoes a transitory actin-driven transient dispersion around the cell nucleus before reassembling near the basal plane. We termed this process Migration Induced Golgi Apparatus Remodeling (MIGAR). During MIGAR, Arp2/3-mediated actin dynamics drives the Golgi dispersion through the MENA-GRASP65 pathway (11). Taken together, our results showed that Golgi-actin association plays an important role in apical-to-basal Golgi relocalization, which is required for maintaining a persistent cell migration. However, once the Golgi apparatus has moved to the basal plane, how its final position is decided remains unknown. In migrating cells, the line connecting the centroid of the cell nucleus to the centroid of the Golgi apparatus defines the cell polarity, which is synonymous with the Golgi polarity. It is, however, not known what factors this polarity depends on. In single cell migration, this polarity axis and the direction of migration show very high co-orientation and are mostly parallel. However, in a migrating epithelial collective, polarity can be perpendicular or even anti-parallel with respect to the global direction of migration. Here, although the cells at the wound edge receive a distinct cue from the cell-free space, and their polarity is mostly parallel to the direction of migration, cells deep within the monolayer are surrounded by neighbors on all sides, which makes the polarity establishment a non-trivial problem.

To understand how the collectively migrating epithelial cells establish their polarity, one can systematically study how different cell biological, geometric, and biophysical factors influence it. Relevantly, previous reports have shown that the kinetic polarity or cell velocity locally orients towards the maximum principal stress, which is known as plithotaxis (2, 12). Hence, the question arises whether the monolayer stress field has any influence on cell polarity. In addition, cell geometry has critical influence on the establishment of cell polarity (13). However, many biophysical and geometric factors are often intricately connected to each other, and it is impossible to perturb one keeping the remaining factors unchanged. To solve this problem, here we propose to use a deep learning framework which can learn the dependency of the cell polarity on any geometric or biophysical factor in an unbiased manner. Importantly, this deep learning framework must be able to process a relational data. Cells in a migrating epithelial monolayer can be modelled as particles interacting with their immediate neighbors, which is local since only neighboring cells interact through cell-cell junctions and is relational since only pairs of cells interact. Hence, a deep learning framework that satisfies this local and relational configuration may be effective in predicting the behavior of collectively migrating epithelial cells. Relevantly graph neural networks (GNNs) have emerged to be useful in the systems where having the knowledge of connections between two entities significantly improves the chance of correct labeling and classification (14–16). Relevantly, a study applying GNN towards understanding glasses showed that GNN could be a powerful tool in predicting the long term dynamics of glassy systems, by taking advantage of the structures hidden in the local neighborhood of the constituent particles. Since, cellular dynamics in the epithelial tissue has several structural and kinetic similarities with the glassy systems (17), we here used GNN towards unbiased understanding of how Golgi polarity gets established in a collectively migrating epithelial cell monolayer. Subsequently, our results elucidated a previously unknown dependency of Golgi polarity on the maximum principal component of the monolayer stress field. Further, this dependency seems to act locally in spite of the global collectivity in the direction of migration.

## RESULTS

### Golgi polarity in collectively migrating epithelial cells

To study the intracellular polarity of the Golgi apparatus in collectively migrating epithelial cells, we cultured a confluent monolayer of Madin-darby canine kidney (MDCK) epithelial cells overnight within the confined area of a culture insert (Ibidi). We then lifted off the confinement to trigger the cell migration. Subsequently, we allowed the cells to migrate for 2, 4, and 6 hours before fixing them with paraformaldehyde. In addition, samples fixed immediately after the confinement lift-off (0 hour) served as the stationary control. We examined the Golgi polarity using antibodies against a Golgi-marker protein, Golgin-97, and calculated the position of Golgi centroid with respect to the centroid of the DAPI-stained cell nucleus (**Figure 1a**). Importantly, our previous work had elucidated that between the apical and basal positioning of Golgi, in stationary and migrating cells respectively, there exists a transitory dispersed state, where Golgi particles are distributed around the nucleus (11). We characterized this dispersed state with a dispersion index (δ) that we had defined previously (11). In this regard, we considered each Golgi vesicle as a point, and measured the distance (*d*) of each point from the Golgi centroid. We then computed the root mean square of the distances and normalized it by the nuclear radius (*R*). A higher δ value (>1.5) represents an equatorially dispersed Golgi, a lower δ value (<1.5) represents a polarized and aggregated Golgi on the basal plane. δ peaked around 1.5-hour post confinement lift-off (**Figure 1b**). Since in this work, we primarily focused on the factors that defined the final Golgi position and polarity, we excluded the cells showing dispersed Golgi and having dispersion index greater than 1.5. Within 2 hours of migration, we found that the cells displayed distinct Golgi polarity (δ<1.5) with respect to the cell nucleus. We broadly categorized the Golgi polarity into three classes, 1: Parallel to the wound direction, 2: Perpendicular to the wound direction, and 3: Antiparallel to the wound direction (**Figure 1a**). This was an important observation considering that previous studies had focused only on the cells at the edge, where the Golgi apparatus predominantly polarize towards the leading edge (18–20). Here we detected that Golgi polarization, especially within the cells that were located away from the wound edge, might not necessarily orient towards the wound edge. Further, we found that this tendency was robust and did not change significantly with the duration of migration (**Figure 1a**, *bar graphs*). This result further suggested that the global direction of migration did not influence the Golgi polarity. Instead, there might be local factors that defined the intracellular polarization of Golgi. To further verify our finding in live cells, we used a modified MDCK cell line that stably expressed Golgi-localized mDsRed fluorescent protein (11). We stained the nuclei of these cells with the Hoechst dye and followed the Golgi reorientation and polarization under a live cell-imaging system (**Figure 1c**). The results from live cell-imaging also confirmed the aforementioned observation (**Figure 1c**), and we found that in some cells the Golgi apparatus polarized towards the wound edge while in significantly high fraction of cells, it localized either perpendicular or antiparallel to the wound edge (**Figure 1c**). Finally, to understand how Golgi polarity depended on cell-cell adhesions, which defined the collectivity in the epithelium, we inhibited the E-cadherin containing cell-cell adhesions by treating the cells with a blocking antibody against E-cadherin, called DECMA-1. In presence of DECMA-1, cells migrated like a single entity, and interestingly, the Golgi polarity always appeared predominantly away from the wound edge and distinct classes of Golgi polarity disappeared (**Figures 1d-e**). Taken together, these results revealed that Golgi polarity in collectively migrating epithelial cells is not defined by overall direction of migration or the direction of wound edge; rather there must exist other factors that define Golgi polarity locally.

**Figure 1:**
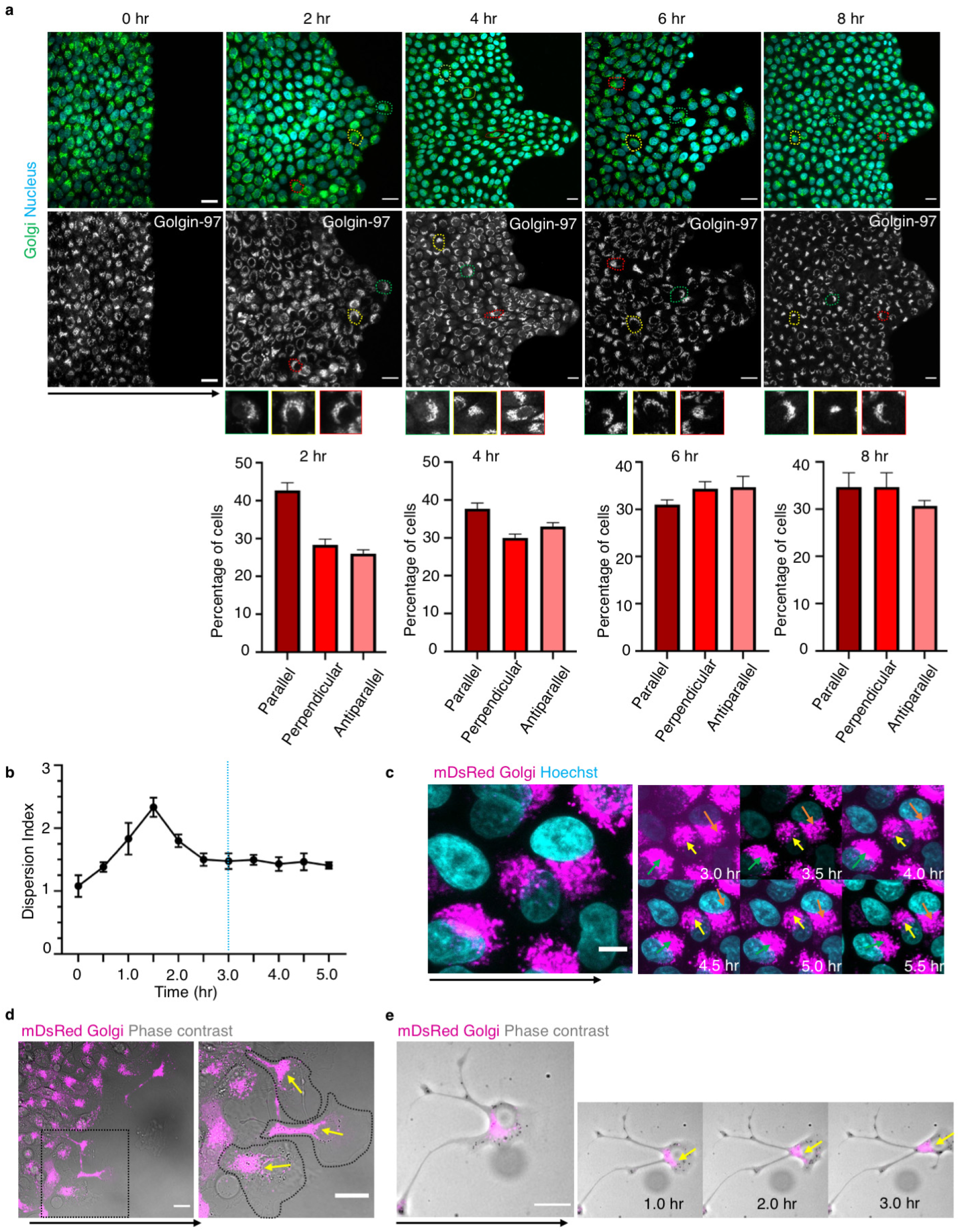
Golgi orientation direction in MDCK monolayer migration does not always align towards the wound. **a)** Immunostained images showing Golgi orientation in cells following 0hr, 2hr, 4hr, 6 hr and 8hr of migration. Highlighted cells show different classes of Golgi orientation with respect to wound, *Green-parallel to wound, Yellow-Perpendicular to wound, Red-Antiparallel to wound*. (*Bottom*) Corresponding quantification in form of bar graphs showing distribution of the three classes of Golgi polarity. Scale bar 20 µm. **b)** The evolution of dispersion index (δ) with time. Dotted line marks threshold after which cells were considered for further experiments. **c)** Live cell time-lapse of mDsRed Golgi expressing cells stained with Hoechst, showing change in Golgi orientation direction for three cells. Golgi alignment with respect to wound, Green-parallel, Yellow-perpendicular, Red-antiparallel. Scale bar 5 µm. **d)** Immunofluorescence images of cell migration carried out in presence of DECMA-1. Cells marked in dotted-black border show Golgi orientation (yellow arrow) aligned away from direction of migration. Solid black arrow denotes direction of migration. Scale bar 20 µm. **e)** Time-lapse montage of a cell migrating in presence of DECMA-1 showing Golgi orientation (yellow arrow) away from direction of migration. Solid black arrow denotes direction of migration. Scale bar 20 µm.

### Geometric and biophysical characterization of collectively migrating epithelial cells

Next, to identify the factors influencing the Golgi polarity, we determined several geometric and biophysical factors associated with individual cells within the migrating collective. Previous studies had shown that the long-range correlation of cellular velocities arose from cells locally orienting their velocity vectors towards the maximum principal stress, by a process known as plithotaxis (12). In addition, collective migration of epithelial cell is also associated with the changes in cell shape. Here, we wondered whether such biophysical or geometric factors might also influence the Golgi polarity. Subsequently we measured cell perimeter (P), area (A), circularity (= P^2^/4πA), shape index (= P/A^0.5^), geometric polarity (orientation of the major axis of the best fitting ellipse), geometric anisotropy (the ratio of major and minor axis of the best fitting ellipse), and the number of vertices as the candidate geometric factors. To this end, we fixed the migrating collectives at 4 hours post confinement lift off and stained the cells for a tight-junction marker protein, ZO-1 (**Figure 2a**, *Left panel*). In addition, we marked the Golgi apparatus with the antibody against Golgin-97 and stained the cell nucleus with DAPI (**Figure 2a**, *Left panel*). ZO-1 formed a thin band at the cell-cell interface and marked the interface more prominently than any other cell-cell adhesion marker proteins. We then used Cellpose platform (https://www.cellpose.org/) (21) to segment out cell boundaries from ZO-1 stained images (**Figure 2a**, *Right panel*). Finally, we fed the output of Cellpose-segmented image **Figure 2a**, *Right panel*) to Tissue Analyzer program (22) to extract the cell identities, which marked the vertices and edges of every cell. We finally processed this output by a custom written software in MATLAB to extract the aforementioned geometric parameters.

**Figure 2:**
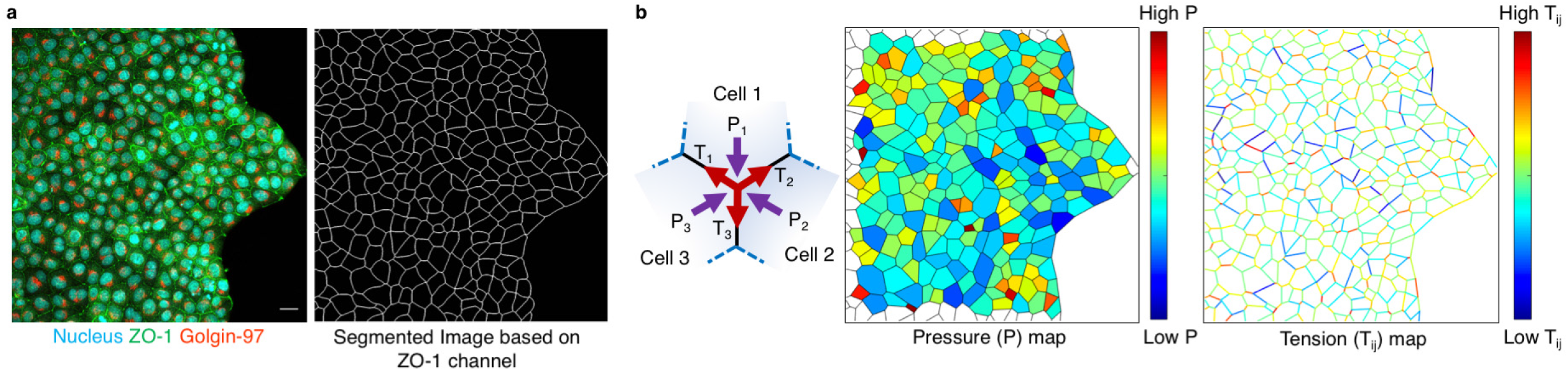
Characterization of geometric and biophysical properties of a migrating epithelial monolayer. **a)** Immunofluorescence image (*Left*) of a monolayer after 4 hour of migration and the corresponding cell segmentation output derived from ZO-1 staining using Cellpose segmentation program. Scale bar 20 µm. **b)** Schematic on left describes the force acting on a three-cell vertex. Cell-cell interface tension (T) and Cell pressure (P). The middle panel shows the relative pressure for each cell, varying between high (red) to low (blue). The right panel shows the relative cell-cell interface tension for each cell, varying between high (red) to low (blue).

We next measured relative tension at cell-cell interface (T_ij_), relative intracellular pressure (P), Batchelor stresses (σ_xx_, σ_yy_, and σ_xy_), principle stress components (σ_1_: maximum, σ_2_: minimum), and Stress anisotropy ([σ_1_ - σ_2_]/[σ_1_ + σ_2_]). To calculate these parameters, we used a previously developed Bayesian force inference method (23, 24). In this method, interfacial tensions (T) and cell pressures (P) were balanced at each vertex. Deriving the force balance equations for all the vertices gave a unified force vector F=Ap, where A (n x m matrix) comprised of coefficients from the force balance equation applied at each vertex, and p was composed of T and P. Assuming that each cell maintains its general shape, the equation is solved for Ap=0. Solving this equation gives an estimation of relative tension and pressure for each cell (**Figure 2b**). However, in case of epithelial sheets, the number of variables is higher than the number of conditions in the equation. To overcome this limitation, a Gaussian function distributed around a constant value was used to get the prior function for tensions. Also, the tensions were assumed to non-negative as cell-cell interface is mostly tensile (24). The estimated interfacial tensions and cell pressures are then integrated to calculate the stress tensor

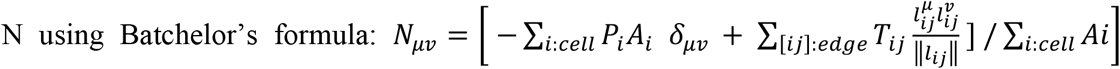

where A_i_ is the area of cell, P_i_ cell pressure, δ is Kronecker’s symbol, T_ij_ is the tension of the edge [ij] between cells i and j, and l_ij_ the vector joining the two vertices of edge [ij]. Batchelor stress can be calculated for individual cell or any region consisting of many cells, based on the desired coarse grain. From this information, we finally calculated the force anisotropy (([σ_1_ - σ_2_]/[σ_1_ + σ_2_]) and the orientation of the maximum principal stress σ_1_ as two candidate biophysical factors. The second factor represented the stress polarity. Again, here we processed the ZO-1 images through Cellpose and Tissue Analyzer. Subsequently, we processed the output of Tissue Analyzer by a custom written software in MATLAB to extract the aforementioned biophysical factors. Finally, we normalized all geometric and biophysical factors so that their value lies between 0 and 1 for non-negative factors and between −1 and 1 for the factors that can assume negative values. Taken together, these geometric and biophysical factors, plus the magnitude of Golgi polarity (the distance between the Golgi centroid and the nuclear centroid) and the distance of the centroid of a cell from the wound edge, constituted the feature vector, which was assigned to each cell. We then asked whether knowing these features will enable us to predict the correct class of Golgi polarity.

### Construction and performance evaluation of GNN

Here, given the collectively migrating epithelial cells assumed a graph-like structure, we reasoned that it could be best modeled by a GNN to predict the Golgi polarity in migrating cells (**Figure 3a**). Under the GNN framework, we imagined that the cells form the nodes of a graph network, where each cell is connected only to its neighboring cells (**Figures 3a-b**). This cell-cell interaction formed the edges of the graph (**Figure 3b**, *middle panel*). The list of normalized geometric and physical parameters then constituted the feature vector, which we assigned to the corresponding node (**Figure 3b**, *middle panel*). Depending on the Golgi polarity of a cell, we also assigned an appropriate class label to the corresponding node, which were as follows: parallel = 1 (blue), perpendicular = 0 (yellow), antiparallel = −1 (red) (**Figure 3b, right panel**). Subsequently, we solved the so called ‘classification problem’ of GNN (25) by passing the feature information from the neighboring nodes to the assigned node, twice (**Figure 3c**). In this regard, we used a special spectral graph convolution method, known as a Chebyshev convolution (ChebConv operator of Pytorch Geometric), which imposes localized convolutional filters via a recursive polynomial decomposition (25). In fact, we tested three other Pytorch Geometric operators including SAGEConv, which implements the GraphSAGE operator (26), GCNConv, which is a graph convolutional operator for semi-supervised classification (27), and GraphConv, which is designed to handle higher-order GNNs (28). However, ChebConv generated the highest accuracy predictions. In our scheme, we convoluted the features of neighboring nodes by ChebConv function twice (**Figure 3c**) and applied the Rectified Linear Unit (ReLU) non-linear activation function in between. The final classification was assigned by passing the output of the second Chebyshev spectral convolution through the LogSoftmax function, which labeled each node to one of the three classes and is advantageous over the Softmax function in terms of enhanced numerical performance and gradient optimization. Once we had defined the network model, we then trained this network with a data for which all node classifications were labeled (**Figure 3d**, *Left panel*). For both training and evaluation run later, we used 20% hidden nodes and 0.5 dropout rates, and ran the program for 600 epochs. Once the network had been training, we tested its performance on a test data (**Figure 3d**, *Right panel*). By comparing the measured Golgi polarity class (**Figure 3d**, *Right panel*) and the GNN-predicted polarity class (**Figure 3d**, *Middle panel*) for all nodes, we achieved approximately 75% accuracy in correctly predicting the Golgi polarity by Chebyshev spectral convolution, which was significantly better than the predictive accuracy of other operators (**Figure 3e**). Taken together, these results showed that collectively migrating epithelial cells could be modeled a graph network (**Figure 3a**) and the Golgi polarity in these cells could be predicted (**Figure 3d**) under the framework of a graph neural network.

**Figure 3:**
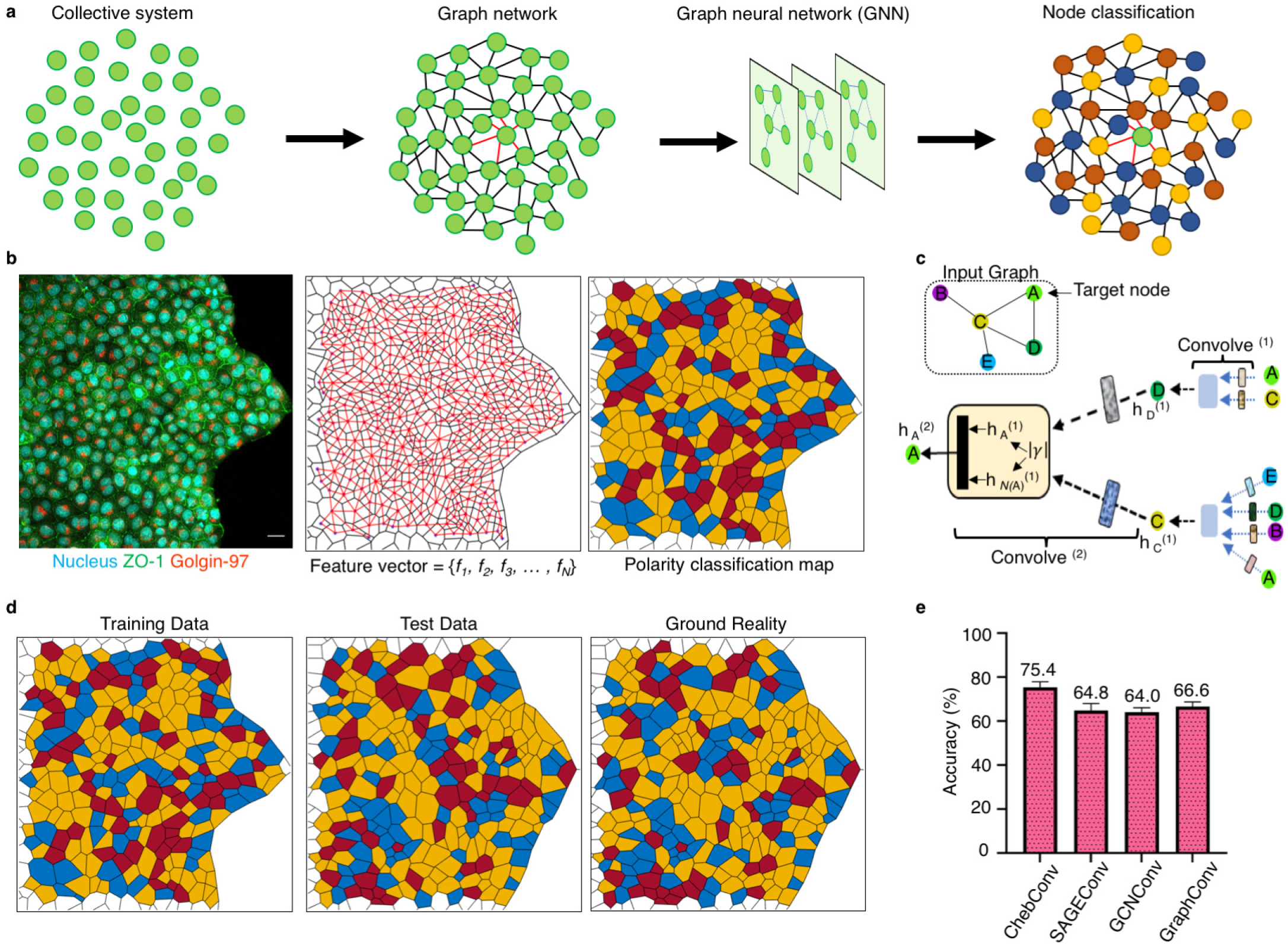
Development of the GNN and performance assessment. **a)** A schematic description of development of the GNN. Each cell of the collective system forms the nodes. Each node connects to the nearest neighboring node to form the Graph network. Nodes are labeled with an appropriate class of Golgi polarity. **b)** The left panel shows the immunostained image used for extracting feature vector of the collective system. Scale bar 20 µm. The middle panel shows the feature vector assigned to each node. The right panel shows the polarity classification map for the collective system. The three different colors represent the three classes of Golgi orientation. Node labels are as follows: parallel = 1 (blue), perpendicular = 0 (yellow), antiparallel = −1 (red) **c)** A schematic of a two layered convolution describing the incorporation of information at one node from its neighboring nodes. Node A receives information from its neighbors, node C and D. In turn, nodes C and D receive input from their neighbors. **d)** A comparison of polarity classification maps obtained from Training data, Test data and Ground reality. **e)** A bar graph comparing the accuracy percentage of the four types convolution method. ChebConv shows the highest accuracy percentage, whereas SAGEConv, GCNConv, and GraphConv show comparable accuracy. The average accuracy percentage shown on top of each bar.

### Correlation between the stress polarity and Golgi polarity

Having predicted Golgi polarity with 75% accuracy, we then asked what could be the most dominating feature that led to this prediction. To answer this question, we took two complementary strategies. First, we studied the average learnt weight of each feature from the first convolution layer, after the network training and testing. In artificial neural network, the neurons gain weight if they are consequential. Our analysis indicated that the weight of the orientation of the maximum principal stress (feature number 10) was significantly higher than the weight of the other features (**Figure 4a**). Secondly, we excluded one feature at a time from our model and studied how it changed the accuracy of the prediction. We subsequently observed the highest drop in prediction accuracy when we excluded the orientation of the maximum principal stress from the feature list (**Figure 4b**). These results suggested that the orientation of the maximum principal stress (stress polarity) predominantly determined the Golgi polarization. In other words, at any point within a collectively migrating epithelial monolayer, polarization of the Golgi apparatus followed the principal direction of the monolayer stress field, showing analogy to plithotaxis (12). We then tested the validity of this GNN-derived prediction by studying the mutual angle (θ) between the Golgi polarity (line connecting the Golgi centroid and the nuclear centroid) and orientation of the maximum principal stress (**Figures 4c-d**). If these two directions did not have any correlation, θ would be uniformly distributed between 0-90º. On the other hand, any correlation between these directions would skew the distribution of θ towards 0º. Plotting the distribution of θ indeed revealed a correlation between the Golgi polarity and orientation of the maximum principal stress (**Figure 4d**). We next tested the locality of this correlation via a progressive coarsening of the stress calculation by increasing the radius of the circle for Batchelor stress calculation (**Figure 4d**). Results obtained from coarsened stress field revealed that correlation between the Golgi polarity and orientation of the maximum principal stress gradually disappeared with increased coarsening, indicating that this correlation is local (**Figure 4d**). Concomitantly, coarsening of the stress field also reduced the predictive accuracy of the GNN model (**Figure 4e**). Taken together, these results indicated that in collectively migrating epithelial cells, the polarization of Golgi apparatus is guided by the principal direction of monolayer stress field.

**Figure 4:**
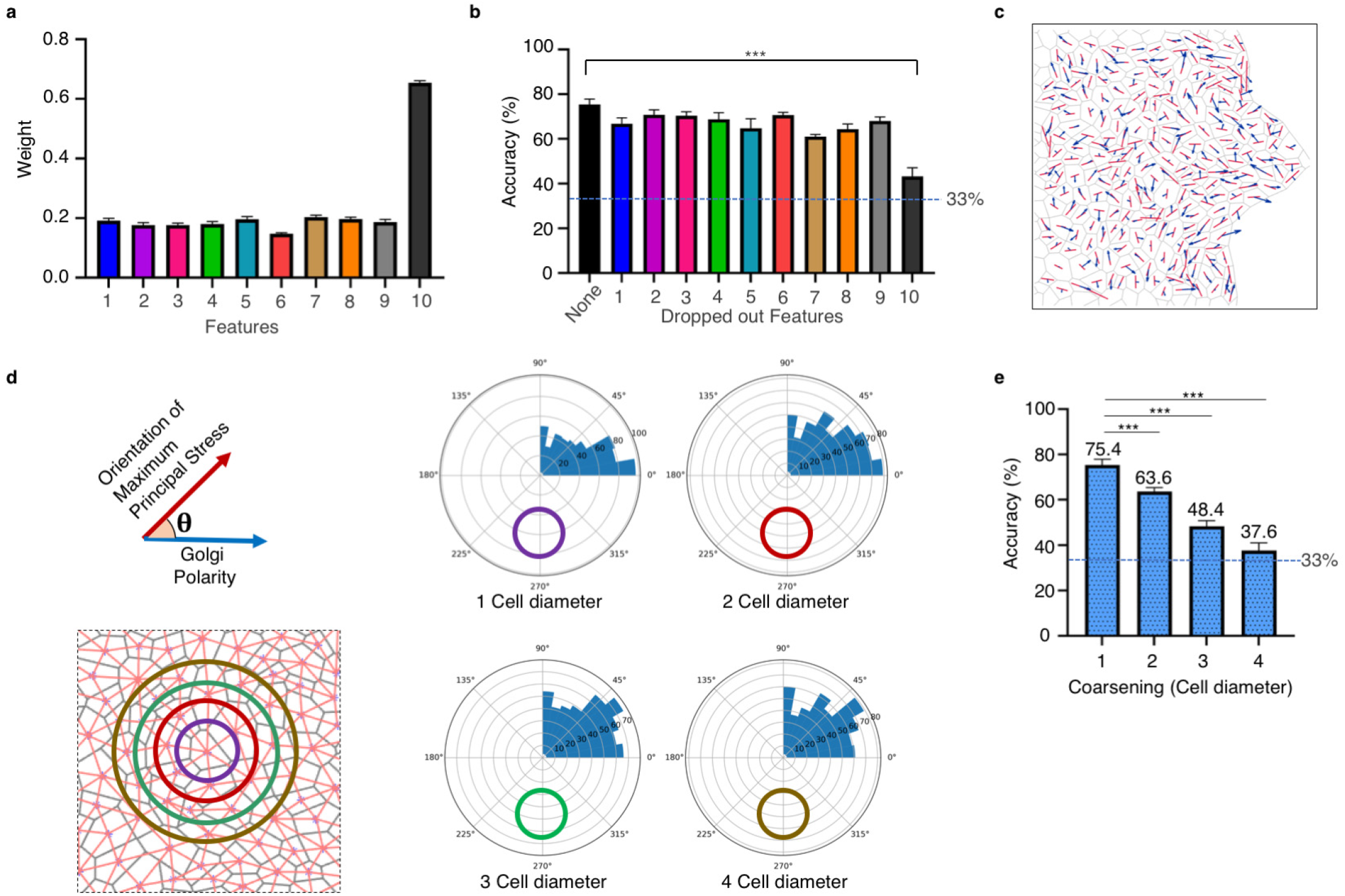
GNN guided discovery of correlation between force-field and Golgi polarization. **a)** A bar graph comparing the weightage of the different parameters. Feature 10 shows the highest weightage. Features: 1-Golgi Polarity Magnitude, 2-Distance from Edge, 3-Cell Area, 4-Circularity, 5-Shape Index, 6-Number of vertices, 7-Geometric Polarity, 8-Geometric Anisotropy, 9-Force Anisotropy, 10-Oreination of Maximum Principal Stress. **b)** Removing the Feature 10-Orientation of Maximum Principal Stress shows a significant drop in prediction accuracy percentage of the model. **c)** Golgi orientation shows a strong correlation with the Orientation of Maximum Principal Stress. **d)** Schematic on left describes the calculation of theta (θ) between Golgi orientation and Orientation of Maximum Principal Stress. The second panel shows the increase in grain size by considering different number of cell diameters. The polar plots show the best correlation between the Golgi orientation and Orientation of Maximum Principal Stress at 1 cell diameter. A gradual decrease (2, 3, and 4 cell diameter) in the correlation occurs with increasing cell diameter. **e)** A bar graph showing the gradual fall in the prediction accuracy of the model with growing coarsening (cell diameter). 1 cell diameter shows the highest prediction accuracy. Average accuracy percentage for each category shown on top of the bar.

## DISCUSSION

The change in Golgi orientation is one of the major reorganization events in the collective migration of epithelial cells. In this context, contrary to the general belief that the direction of Golgi polarization should be predominantly be parallel to the direction of migration or the wound edge, we found that the Golgi polarization comes in a varied range of classes, including parallel, perpendicular, and antiparallel orientation with respect to the wound direction. In our quest to understand this variation, we developed a graph neural network-based deep learning tool to understand and computed various factors of a cell collective, including geometric factors and biophysical factors. This tool allowed us to build a strong information set for a cell collective and enabled us to isolate the most important factor that affects the Golgi polarity within a collectively migrating epithelial monolayer. Here we found that the orientation of the maximum principal stress or the stress polarity showed the strongest correlation with the Golgi polarity. Further, this correlation between the stress and Golgi polarity strongly depends on the immediate local environment of individual cells. While from our study, GNN emerged to be a very useful tool in understanding the Golgi polarity in a migrating collective, we believe that this effort is the just the beginning, and the true power of GNN must be exploited to understand the collective cell behavior in greater details. Possibly a larger volume of information is needed to add another dimension to the understanding of collective cell migration by GNN.

Although our GNN includes several factors, as described above, it is not an exhaustive list of factors. The addition of other factors such as the polarity of other organelles (including centrosome and lysosome), orientation of cytoskeletal elements, and the energy state of the cell could be advantageous. Further, future works should focus on using GNN to understand collective cell migrations in model organisms to elucidate the underlying principles of tissue remodeling and morphogenesis. Pertaining to detailed structure of GNN, one should note that while we have tested the performance of four distinct convolution operators, namely ChebConv, SAGEConv, GCNConv, and GraphConv, there are other operators in the Pytorch Geometric database whose performance remain to be tested in this context. Another important component that can be optimized is the non-linear activation function. Here, we have used ReLU, which is the most commonly used activation function because of its simplicity. But the effect of other non-linear functions on the prediction accuracy needs to be tested. Thus far we have managed to achieve nearly 75% accuracy, which, given the complexity of the problem, is a very good performance against a random accuracy of 33%. Although further optimization of the GNN architecture should improve this accuracy, we do not expect that the main conclusion of the work, namely the stress polarity-Golgi polarity correlation, to be qualitatively affected by any further improvement in the prediction accuracy.

While the current work shows an interesting aspect of Golgi orientation in a migrating collective, it is limited by the lack of a satisfactory molecular explanation. How the monolayer stress polarity information is transduced to Golgi polarity remains unanswered. It is likely that the actin cytoskeleton, one of the major force-responsive structures, is facilitating the communication between the two. In fact, an existing model describing the role of PtdIns (4)P/GOLPH3/MYO18A pathway might explain our observations (29). In migrating cells, the F-actin distribution shows a front-to-rear gradient, with a higher concentration at the front and a lower concentration at the rear (30–33). The increased F-actin at one end of the cell possibly leads to an increased interaction with myosin II and other non-muscle myosins. This increased force-bearing through the actomyosin complex might create a force imbalance leading to Golgi reorientation. Consequently, the Golgi reorientation might occur in a direction where this force imbalance resolves. Further studies need to be performed to test this hypothesis. Nevertheless, in this work, the unique usage of GNN in collective cell migration provides the first step towards such endeavor. Just as the discovery of plithotaxis (12) led to the discovery of the merlin-mediated molecular mechanotransduction pathway, which governs the long-range correlation of cellular motions (2), we envisage that our discovery of the stress polarity-Golgi polarity correlation will lead to the discovery of a signaling pathway linking the physical cues to organelle reorganization.

## ACKNOWLEDGMENTS

We thank the Collective Cellular Dynamics (CCD) laboratory members for critical discussion. We sincerely thank Raphaël Clément for the source code of Bayesian force inference program. T.D. is a Department of Biotechnology (DBT)/Wellcome Trust India Alliance intermediate fellow and partner group leader of the Max Planck Society, Germany. The authors sincerely acknowledge generous funding from a partner group grant from the Max Planck Society, Science & Engineering Research Board (SERB), DST, Govt. of India (Sanction Order No. CRG/2021/003907), and intramural funds at Tata Institute of Fundamental Research, Hyderabad from the Department of Atomic Energy Energy, India (under Project Identification No. RTI 4007). Authors also acknowledge the generous funding by the Human Frontier Science Program.

## MATERIALS AND METHODS

### Cell culture

All the experiments were done using the Madin-Darby canine kidney (MDCK) epithelial cell line. Tetracycline-resistant Wild-type MDCK (MDCK-WT) cell lines were a gift from Yasuyuki Fujita. Cells were cultured in Dulbecco’s modified Eagle’s medium supplemented with GlutaMax (Gibco) with 10% fetal bovine serum (tetracycline-free FBS, Takara Bio) and 10 U mL-1 penicillin and 10 µg mL-1 streptomycin (Pen-Strep, Invitrogen) in an incubator maintained at 37°C and 5% CO2, unless mentioned otherwise.

### Immunofluorescence

Cell fixation was done with 4% formaldehyde diluted in 1x phosphate-buffered saline (PBS, pH 7.4) at RT for 10 minutes, followed by 1X PBS washes (three times). Cell permeabilization was carried out with 0.25% (v/v) Triton X-100 (Sigma) in PBS for 10 min at RT followed by washing thrice with PBS to remove the reagent. To block non-specific antibody binding samples were incubated in 2% BSA in PBST (0.1% v/v Triton X-100 in 1X PBS) at RT for 45 minutes. The blocking buffer was removed after 45 minutes, and the primary antibody dilution prepared in blocking buffer was added to the samples for 60 minutes at RT or at 4°C overnight. Afterward, samples were washed twice with PBST and thrice with 1X PBS. Next, secondary antibody tagged with AF 488/594 were (same dilution as primary) prepared in blocking buffer and added to the sample for 60 minutes at RT. To counterstain cell nuclei and F-Actin, the samples were added with a DNA-binding dye 4′,6-diamidino-2-phenylindole (DAPI, 1 µg mL-1 in PBS, Invitrogen) and phalloidin conjugated Alexa Fluor dye (Invitrogen) along with the secondary antibody solution. Lastly, a thorough washing of the samples was done with PBST and PBS before imaging. Golgin-97 (CST), GRASP65 (Thermo Fisher), ZO-1 (CST), DECMA1 (Sigma). All antibodies were used in 1:200 dilution.

### Calcium switch experiments

To block E-cadherin mediated AJs, confluent cell monolayers grown in culture inserts were first washed twice with calcium–magnesium free HEPES and then incubated with 4 mM EGTA (Sigma) in HEPES buffer (Invitrogen, pH 7.4) for 30 min at 37°C in a 5% CO2 humidified incubator. Following the incubation periods, the calcium free medium was replaced with the cell culture medium with or without E-cadherin blocking monoclonal antibody DECMA-1 (Millipore, 10 µg mL−1), and cells migration was carried out for 4 hours at 37°C in a 5% CO2 humidified incubator.

### Statistical analysis

All statistical analysis was done using GraphPad Prism. Two-tailed unpaired t-test was used to quantify the effect of dropping different features and coarsening, on accuracy percentage.

### Computation of geometric and biophysical parameters

Relative pressure within each cell and relative tensions at cell-cell interface were determined from the cell geometry using a Bayesian force inference technique (23, 24). To this end, DIC, phase-contrast, or fluorescent images marking cell-cell interface was segmented using a Deep learning-based program, Cellpose (21). Output of this program was then processed by Tissue analyser (22) to assign cell vertices and edges. The list generated by Tissue analyser was then fed into a program written in MATLAB (MathWorks), which implemented Bayesian force inference. In this method, a matrix describing balance of interfacial tensions and intracellular pressures at each vertex (Fig. 1c, *bottom*) was derived. To solve this underdetermined linear system, again Bayesian inversion was implemented and all variables and parameters of the problem were determined with certain probability. Details of the probabilistic determination had been described in detail elsewhere (23). Both relative pressure and tension, obtained by Bayesian force inference, are unitless quantity. Systematic comparisons of force inference with laser ablation experiments recently showed the validity of this method in diverse range of tissues (24).

### Implementation of graph neural network (GNN)

The graph neural network was written in Python programing language on Anaconda distribution. To implement the GNN, PyTorch Geometric (https://pytorch-geometric.readthedocs.io/en/latest/) library was used. PyTorch Geometric is built upon PyTorch to facilitate writing and training GNNs for a vast range of applications pertaining to structured data. We used the Chebyshev spectral graph convolutional operator with in_channels = 10, out_channels = 20, K=2, and normalization = symmetric in the first convolutional layer and with in_channels = 20, out_channels = 3, K=2, and normalization = symmetric in the convolutional layer. ReLU and LogSoftmax were chosen as the non-linear activation function and classification, respectively. It was observed that lesser number of hidden units led to slower convergence, higher loss, but better accuracy. The number of hidden layer was fixed at 20, which is double the number of features. Dropout rate was fixed at 0.5. For accuracy calculation, we discounted the cells with dispersed Golgi (δ>1.5).

